# Foxi3 Suppresses Signaling Center Fate and is Necessary for the Early Development of Mouse Teeth

**DOI:** 10.1101/2022.07.18.500404

**Authors:** Isabel Mogollón, Niko Kangasniemi, Jacqueline Emmanuel Moustakas-Verho, Laura Ahtiainen

**Author notes:** Cell and Tissue Dynamics Research Program, Institute of Biotechnology/Helsinki Institute of Life Science, P.O. Box 56 (Viikinkaari 5), FIN-00014 University of Helsinki, Finland.

## Abstract

Tooth morphogenesis is regulated by ectodermal-mesenchymal interactions which are controlled by reiteratively used, evolutionarily conserved pathways. How these signals translate into different cellular behaviors is beginning to be understood. Embryonic cell behaviors are regulated by signaling centers that define organ position, size, and shape. The recently identified initiation knots (IKs) and the well-known enamel knots (EK) are tooth epithelial signaling centers that drive proliferation of the neighboring cells and are involved at different stages of morphogenesis, from the early epithelial thickening to the final formation of shape. Signaling center cell fate and maintenance can be regulated by numerous factors. Here, we studied the early stages of murine incisor and molar epithelial morphogenesis and overcame the previous shortage of early-stage mutant models to functionally manipulate the initiation processes of tooth morphogenesis. We achieved the early conditional knock down of the forkhead-box transcription factor Foxi3 during tooth initiation and used imaging approaches to explore cellular and molecular disease mechanisms, specifically those related to signaling center dysfunction in tooth dysplasia. We show that early deficiency of Foxi3 in incisors and molars frequently arrests growth at bud stage, whereas later knockdown of Foxi3 affects tooth downgrowth and shape. Cell-level analyses revealed a decrease in epithelial proliferation and the ectopic appearance of cells with hallmarks of signaling centers: quiescent cell state and canonical Wnt activity. However, the distribution of these cells was wider all over the tooth buds with abnormal decrease of apoptosis. We postulate that, depending on *Foxi3* expression levels, the bud cells shift commitment into signaling center fate, ultimately leading to growth arrest or growth/shape changes, implicating Foxi3 as a regulator of cell fates between the signaling centers and proliferating bud cells.

## Introduction

Ectodermal organs, such as hair and teeth, develop through embryonic epithelial-mesenchymal interactions across similar stages that have been defined morphologically and genetically. Tooth development begins with the initiation stage, which comprises a thickening of the epithelium (placode), followed by the bud stage with a condensation of the mesenchyme and epithelial invagination (Pispa and Thesleff, 2003, Jernvall and Thesleff, 2000). These morphogenetic events are controlled by well-known conserved pathways that are used reiteratively, but how these actions translate to different cellular behaviors is only beginning to be understood (Ahtiainen et al., 2014, Kuony and Michon, 2017, Biggs et al., 2018, Mogollón et al., 2021, Trela et al., 2021). Specialized relatively small groups of cells called signaling centers play an essential role regulating cell behaviors, doing so by secreting factors such as sonic hedgehog (Shh), and molecules belonging to the Wnt, Fgf, and Bmp families. In tooth, these signaling centers form sequentially and control tissue remodeling to define organ position, size and shape (Jernvall and Thesleff, 2012, Mogollón et al., 2021).

The best characterized signaling centers in tooth development are the enamel knots (EK) forming at the tip of the epithelial tooth bud followed by the formation of secondary enamel knots (sEK) at the tips of future cusps in the molar (Jernvall and Thesleff, 2000). The early development of both incisors and molars in mice is regulated by an earlier forming epithelial signaling center, the initiation knot (IK; Ahtiainen et al., 2016; Mogollón et al., 2021). IK cells exit the cell cycle (G1/G0) at tooth placode stage, further condense via cell migration forming the mature signaling center, and drive proliferation of the neighboring dental epithelial cells in the growing bud. Subsequently, the IK remains quiescent and then is apoptotically silenced as the primary EK arises independently (Mogollón et al., 2021). IK maturation involves the interaction of cells with high canonical Wnt activity and the non-proliferative Shh-expressing cells (Mogollón et al., 2021). Signaling center cell fate and maintenance can be regulated by several factors such as ectodysplasin/NFκB (Eda); while a novel regulator, transcription factor Forkhead box I3 (Foxi3), likely maintains the proliferating population (Jussila et al., 2015).

Foxi3 is an important regulator of ectodermal development, belonging to the superfamily of Forkhead box transcription factors which include molecules with diverse functions, such as chromatin modifiers, activators and repressors (Benayoun et al., 2011, Lam et al., 2013). Foxi3 expression locates in the epithelium of several ectodermal organs (Shirokova et al., 2013); in tooth, it is expressed in different cell populations at several developmental stages (Shirokova et al., 2013;Jussila et al., 2015). A mutation of Foxi3 found in hairless dogs causes missing and abnormally shaped teeth (Drögemüller et al., 2008). This phenotype resembles hypohidrotic ectodermal dysplasia (HED) in humans, caused by mutations in the Eda gene that is a regulator upstream of Foxi3 (Shirokova et al., 2013). In mice, homozygous mutation is lethal due to defects in branchial arch formation (Edlund et al., 2014). To study the role of Foxi3 during ectodermal organ morphogenesis, Foxi3 conditional knockouts have been produced (Foxi3cKO). A conditional bud stage deletion of Foxi3 targeting epithelial expression with the keratin-14 promoter results in a fusion of the molar teeth (Jussila et al., 2015). This bud stage Foxi3 mouse mutant shows a milder phenotype compared to the one seen in dogs (which lack some teeth), this might be because in mice the absence of Foxi3 is conditional and occurs later in development (Kupczik et al., 2017). This model shows efficient conditional knock down of Foxi3 from embryonic day E13, making it a valuable tool to study the interaction between budding and tooth shape. Nevertheless, Foxi3 is expressed already at placode stage (∼E11), so the study did not account for the possible effects caused by knocking out Foxi3 during tooth initiation and IK regulation.

Many genetic defects causing tooth agenesis syndromes in humans have equivalent mouse models, and the foremost defect is already seen at the initial stages of morphogenesis (Tummers and Thesleff, 2009; Hallikas et al., 2021). However, because of a shortage of early-stage models, mutants have been mainly explored at bud stage and beyond. In addition, the use of conventional two-dimensional sectional imaging methods has found it difficult to visualize the small dimensions of morphogenetic events during tooth initiation (Mogollón and Ahtiainen, 2020). Recent improvements in imaging methods have brought a better understanding of the cellular mechanisms of very early tooth morphogenesis that have been elusive for decades due to the lack of realistic three-dimensional datasets (Sharir and Klein, 2016). Foxi3 is expressed in different patterns in the placode and later in bud stage, so it is likely that it has distinct roles depending on the context. Thus, a mechanistic understanding of early developmental events in teeth is needed and studying Foxi3 offers an opportunity to do so. For these reasons, we generated an early stage Foxi3 conditional mouse model and used imaging approaches to explore cellular and molecular disease mechanisms, specifically those related to signaling center dysfunction in tooth dysplasia mouse models lacking Foxi3.

We used confocal fluorescence whole-mount tissue imaging to visualize the bud stage K14-Foxi3cKO model and established an efficient early conditional Foxi3 knockdown model using the inducible Sox2^CreERT2^. This Cre driver is specifically active in cell populations that express Foxi3 from E11 (Ahtiainen et al., 2016, current study). Here we show that Sox2 ^CreERT2^-Foxi3cKO incisors and molars are very frequently arrested at the bud stage, unlike K14^Cre43^-Foxi3cKO teeth which develop into mature molars but display flat and abnormal shapes (Jussila et al., 2015). Both Foxi3 mutants displayed defects in apoptosis, and the normally proliferating bud cells exited the cell cycle and abnormally expressed signaling center markers. Thus, we postulate that the bud cells shift commitment into signaling center fate, most prevalently in the early stage. Such changes in signaling center fate definition, extended cell cycle exit, and hampered silencing, suggest a novel cellular disease mechanism in ectodermal dysplasia. We propose that the function of Foxi3 in the initial stages of tooth development lies in the regulation of cell fates between the signaling centers and proliferating bud cells, and depending on Foxi3 expression levels, ultimately leading to growth arrest or growth/shape changes.

## Results

### The Sox2^CreERT2^Foxi3cKO is a novel mouse model for the early epithelium knock down of Foxi3 in teeth

Efficient conditional knock down of Foxi3 in the epithelium can be achieved from embryonic stage E13 (tooth bud stage) with the Keratin 14 Cre mouse (K14^Cre43^; Foxi3 ^−/floxed^; K14^Cre43^FoxicKO mouse). However, Foxi3 is expressed already earlier at the dental lamina stage, suggesting that this transcription factor might have a function in tooth placode and bud formation from E11 (Shirokova et al., 2013). We established an early stage conditional Foxi3 knockdown mouse model using the Sox2^CreERT2^ inducible driver, in a mixed ICR and NMRI background (Sox2^CreERT2^; Foxi3^+/−^; hereafter Sox2^CreERT2^Foxi3cKO mouse). We chose this model since Sox2 and Foxi3 are expressed in the same cell populations at early stages of tooth morphogenesis (Ahtiainen et al., 2016) and it allows a very specific time control for conditional Foxi3 knock down (Figure 1a). To test the efficiency of Foxi3 knock down in Sox2^CreERT2^; Foxi3^+/−^ mice, Foxi3^floxed/floxed^ females were crossed with Sox2^CreERT2^; Foxi3^+/−^ males and tamoxifen was administered twice to pregnant females at E10.5 and E11.5 (Figure 1b). We also analyzed the efficiency of the Cre-driver used with qRT-PCR and confirmed the downregulation of Foxi3 expression in both mutant models: in Sox2^CreERT2^Foxi3cKO teeth at E12.5 and in K14^Cre43^Foxi3cKO teeth at E13.5 (Figure 1c). To complement our analysis, we tested the induction efficiency of the Sox2^CreERT2^ inducible Cre driver, ROSA-tDtomato fluorescent inducible reporter females were crossed with Sox2^CreERT2^ males, and tamoxifen was injected on pregnant females at embryonic day 10 (Figure 1d). We observed Cre-induction in the same cell populations that express Foxi3 at E11 and E12, within the epithelial dental placode (Figure 1d). For analyses of cellular behaviors, we crossed parental animals (Foxi3^floxed/floxed^ females and Sox2^CreERT2^; Foxi3^+/−^ or K14^Cre43^; Foxi3^+/−^ males) carrying other reporter lines. The Foxi3^+/floxed^ animals were used as littermate controls.

**Figure 1.**
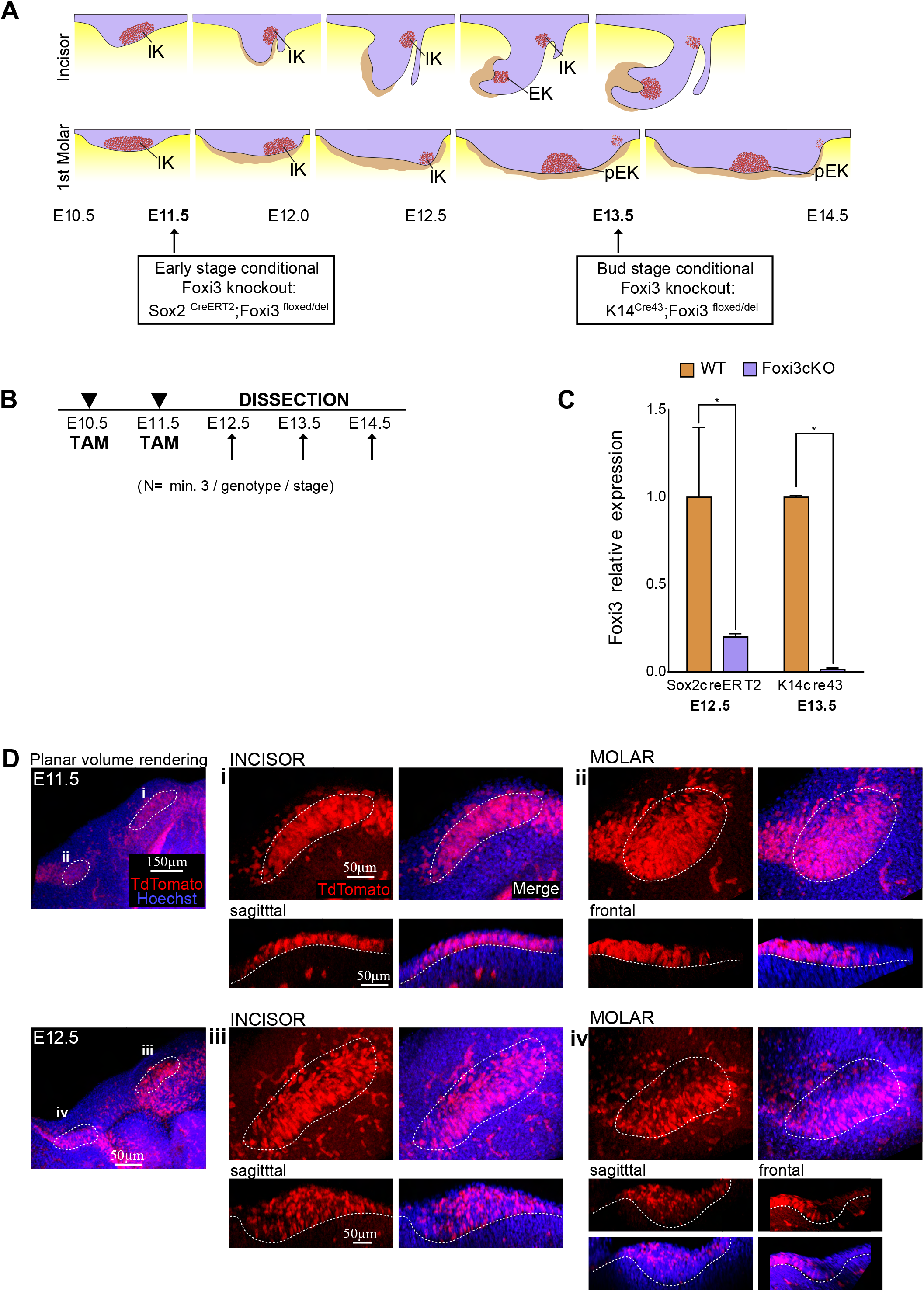
Novel mouse model for early knock down of Foxi3 in teeth. (A) Experimental setup for Cre induction with tamoxifen injections. The Sox2^CreERT2^ inducible Cre driver was used to achieve conditional knock down of Foxi3 in the dental epithelium in a time-specific manner. Pregnant females were injected twice by intraperitoneal injections with tamoxifen at E10.5 and E11.5 (daily dose: 2.5mg/30g weight per mouse). (B) Relative expression of *Foxi3* analyzed by qRT-PCR in embryonic mouse teeth. Foxi3 gene expression in E12.5 Sox2^CreERT2^Foxi3cKO (n=3) and E13.5 K14^Cre43^ (n=3) mutant teeth were compared to wild type tissue of corresponding litters (n=3+3). Foxi3 expression is decreased in Sox2^CreERT2^Foxi2cKO E12.5 mutant teeth by 0.2-fold, and in K14^Cre43^Foxi3 E13.5 mutant teeth by 0.02-fold. Gene expression levels were normalized to GAPDH expression. Results shown as mean±s.d. Statistical significance of the difference between mutant and control samples was tested with Mann-Whitney *U*-test. *P*-value of 0.05 was chosen as the significance threshold. (C) ROSA-tdTomato fluorescent reporter showing the sites of Sox2^CreERT2^ activity at different time points. ROSA-tdTomato fluorescent reporter females, which display a red fluorescent cytoplasmic protein in the presence of the Cre protein, were crossed with Sox2^CreERT2/+^ males in order to detect the level of efficiency of this model to label Sox2-expressing cells. Pregnant ROSA-tdTomato females were injected with tamoxifen one day before dissection (at E10.5 or E11.5) and dissected at E11.5 or E12.5, respectively. Confocal microscopy immunofluorescent whole-mount images of Sox2^CreERT2^; R26R^tdTomato^ mandibles at E11.5 and E12.5 stages. Immunolabeled with ROSA-tdTomato label (red), and nuclei stained with Hoechst 3342 (blue). (Planar view) Each figure is a volume rendering of the xy plane. (Sagittal and frontal view) Each figure is an optical section in the z plane. When labelling was induced at E10.5 and analyzed at E11.5 most incisor and molar cells were tdTomato+, indicating that the tamoxifen induction causes high labelling efficiency at this dosage and timing. Likewise, after tamoxifen administration at E11.5 and analysis at E12.5, high number of cells were also tdTomato+.

### Early knockdown of Foxi3 causes bud growth defects in incisors and molars

In order to study the cellular mechanisms during early tooth morphogenesis in WT and Sox2^CreERT2^Foxi3cKO mutants, we studied cell cycle dynamics on epithelial bud invagination in incisors and molars from mouse mandibles at placode stage (E12.5), bud stage (E13.5) and early cap stage (E14.5). The epithelium of ectodermal organs can be visualized with the transgenic reporter mouse expressing Keratin17-GFP (K17-GFP) (McGowan and Coulombe, 1998). We used 3D surface renderings of the K17-GFP reporter to visualize the bud invagination process, and size and shape changes. In all analyzed stages (E12.5-E14.5), K17-GFP expression was present in both littermate WT controls and Sox2^CreERT2^Foxi3cKO mutant teeth. At E12.5, WT placodes started to invaginate to become a bud, whereas mutant incisors and molars were less invaginated (Figure 2a). At E13.5, WT teeth were invaginated and started to go through early cap stage, while mutant teeth were flatter and smaller (Figure 2b). At E14.5, WT incisors and molars already showed a cap shape, but mutant teeth were arrested at bud stage and much smaller (Figure 2b). Quantifications of bud volume showed that mutant buds were significantly smaller already at E12.5 and E13.5 (Figure 2c). Notably, we observed variation in the severity of tooth phenotype in the mutants, so we speculate that depending on the efficiency of tamoxifen induction (and therefore Foxi3 knock down) different phenotypes are present in E14.5 mutant teeth. Most clearly classified in incisors, teeth displayed three different phenotypes: completely arrested small bud, bigger bud, and smaller / not fully mature cap (Figure S1a). We scored these three different phenotypes in mutant incisors and measured tooth size, observing a decrease in bud length compared to the WT phenotype depending on the severity of Foxi3 knock down (Figure S1b). Estimation of the total number of individuals per phenotype showed that 75% of Sox2^CreERT2^ mutant individuals developed growth arrest at bud stage. This phenotypic continuity suggests that even a partial knockdown of Foxi3 expression in dental epithelia at placode stage is sufficient to induce an impairment in growth, and if the deficiency is extended enough, even cause a developmental arrest. We therefore conclude that early conditional knock down of Foxi3 in the Sox2^CreERT2^Foxi3cKO mutant model causes a decrease in tooth size in both incisors and molars, and when the Foxi3 knockdown is efficient enough, it causes growth arrest at bud stage.

**Figure 2.**
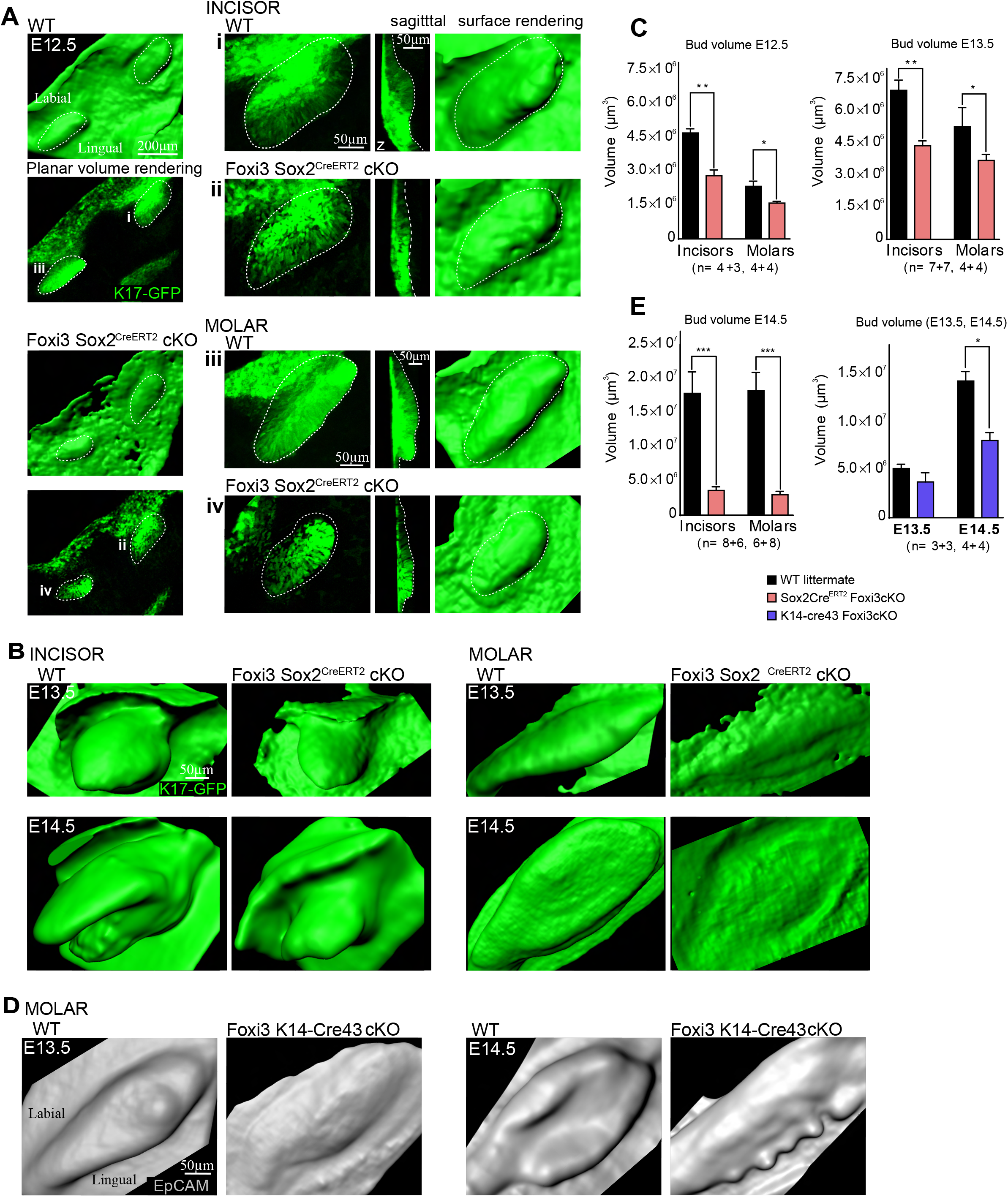
Early stage knock out of Foxi3 arrests development at bud stage, knock out of Foxi3 at later stages affects tooth shape. Bud invagination was studied in early stage Sox2^CreERT2^ and bud stage K14^Cre43^ Foxi3 conditional knockout mutants. Sox2^CreERT2^; Foxi3^del/wt^ animals were injected with tamoxifen at E10 and 11 and mandibles dissected and fixed at E12.5-E14.5 in order to achieve Foxi3 knock down at E11. (A) Whole-mount confocal fluorescence images incisors and molar epithelial buds with keratin 17-GFP (green). (Planar view) Each figure is a volume rendering of the xy plane. (Sagittal view) Each figure is an optical section in z plane of wt and Sox2^CreERT2^Foxi3cKO mutant teeth. Tooth epithelium borders are delineated with a dotted line. (A) K17-GFP reporter (green) reporter was used to visualize the forming tooth buds in early stage Sox2^CreERT2^Foxi3cKO mutants. The mutants showed a clear difference compared to WT from E12.5, the mutants displayed smaller and flatter buds compared to WT tooth buds. (B) Similarly, at E13.5, Foxi3cKO mutant buds are flatter and overall smaller than the WT counterparts. At E14.5 mutant bud growth is arrested and does not progress to cap stage. (C) Quantifications of bud volume show that from E12.5 to E14.5 Sox2^CreERT2^Foxi3cKO mutant incisors and molars are significantly smaller (N_E12.5_=4+3, 4+4; N_E13.5_=7+7, 4+4; N_E14.5_=8+6, 6+8). (D) Whole-mount confocal fluorescence images of molar epithelial buds of Foxi3 K14^Cre43^ bud stage deletion model with the epithelium visualized with immunofluorescence staining (EpCam, grey). Bud stage K14^Cre43^Foxi3cKO mutants showed a clear difference in E13.5 and E14.5 molar phenotype when compared with WT molars. EpCam surface renderings show a smaller and flatter mutant molar shape at E13.5, and E14.5 mutant molars display abnormal epithelial protrusions. (E) Quantifications of molar bud volume of the bud stage K14^Cre43^Foxi3cKO mutant model show that from E13.5 to E14.5 mutant molars are smaller (N_E13.5, E14.5_=3+3, 4+4). However, the bud volume differences are less pronounced compared to those observed with the early stage Sox2^CreERT2^Foxi3cKO mutant model. For all statistical tests: mean±sem, nonparametric Mann-Whitney *U*-test, p<0,05 *, p<0,01**, p≤0.001***.

We also studied bud invagination in the bud stage K14^Cre43^ Foxi3 conditional knockout mutant (Figure 2d-e). We studied molars at E13.5 and E14.5, when there is a clear difference in phenotype of the bud stage K14^Cre43^Foxi3 conditional knockout mutants compared to WT molars (Jussila et al., 2015). The pan-epithelial marker EpCam immunofluorescence staining was used to visualize the epithelium, and molar bud 3D surface renderings of EpCam at E13.5 showed that K14^Cre43^Foxi3cKO mutant molars were flatter (Figure 2d). At E14.5, mutant molars displayed an abnormal shape: the mutant molar epithelium showed extra epithelial foldings at the lingual side (Figure 2d). Quantifications of bud volume showed that mutant buds start to be smaller at E13.5, and the size differences between WT and mutant molars become significant at E14.5 (Figure 2e).

### Growth defect caused by cell cycle exit increase in Foxi3cKO mutant teeth

Signaling centers in ectodermal placodes commonly go through cell cycle exit (Ahtiainen et al., 2014; Ahtiainen et al., 2016; Mogollón et al. 2021). The Fucci cell cycle indicator mouse reporter can be used to identify cell cycle phases and a surrogate marker for signaling centers in early developing incisors and molars. Fucci mice express nuclear red in G_1_/G_0_ phase (Cdt1-mKO) (Sakaue-Sawano et al., 2008). We used confocal fluorescence microscopy of Fucci whole-mount mandibles to study the G_1_/G_0_ signaling centers in developing WT and Foxi3cKO mutant incisors and molars, stages E12.5-E14.5 (Figure 3). At E12.5, WT incisors and molars showed Fucci G_1_/G_0_ initiation knot (IK) cells located mesially in the developing bud, as previously published (Figure 3a; Ahtiainen et al., 2016; Mogollón et al. 2021). At E13.5, WT incisor and molar IKs remained in the mesial part of the tooth buds, close to the epithelial surface, as the emerging EKs appeared deeper in the bud (Figure 3a). On the other hand, at E12.5 Sox2^CreERT2^ Foxi3cKO mutant incisors and molars displayed expanded IK area, and at E13.5 mutant teeth also showed a more scattered distribution of Fucci G_1_/G_0_ cells in not only the signaling center areas but also all over the bud, where the normally proliferating cell population resides in WT teeth (Figure 3a; Ahtiainen et al., 2016; Mogollón et al. 2021). By E14.5, WT teeth were in matured cap stage with well-formed EKs, as compared to mutant teeth in which growth was arrested in late bud stage, probably due to the high number of non-proliferative cells (Figure 3a). Quantification of G_1_/G_0_ cells at E12.5 and E13.5 stages indicated a higher number of total Fucci G_1_/G_0_ positive cells in mutant incisors and molars (Figure 3b). Altogether, mutant teeth show a higher number and scattered distribution of Fucci G_1_/G_0_ positive cells instead of being confined to the IK and EK signaling center regions. The mutant IKs were bigger, and more bud cells exit the cell cycle in mutant incisors and molars at bud stage. This data confirmed that the absence of the Foxi3 transcription factor correlates with an increase of cell cycle exit throughout the whole incisor and molar teeth; subsequently, this proliferation imbalance causes the disruption of proper EK formation. The growth arrest observed in mutant teeth (Figure 2) might be due to an increase of cell number in the non-proliferative epithelial population already at E12.5 (Figure 3a). The Sox2^CreERT2^; Foxi3^+/−^ mouse is, therefore, a successful model to study early signaling center dysregulation.

**Figure 3.**
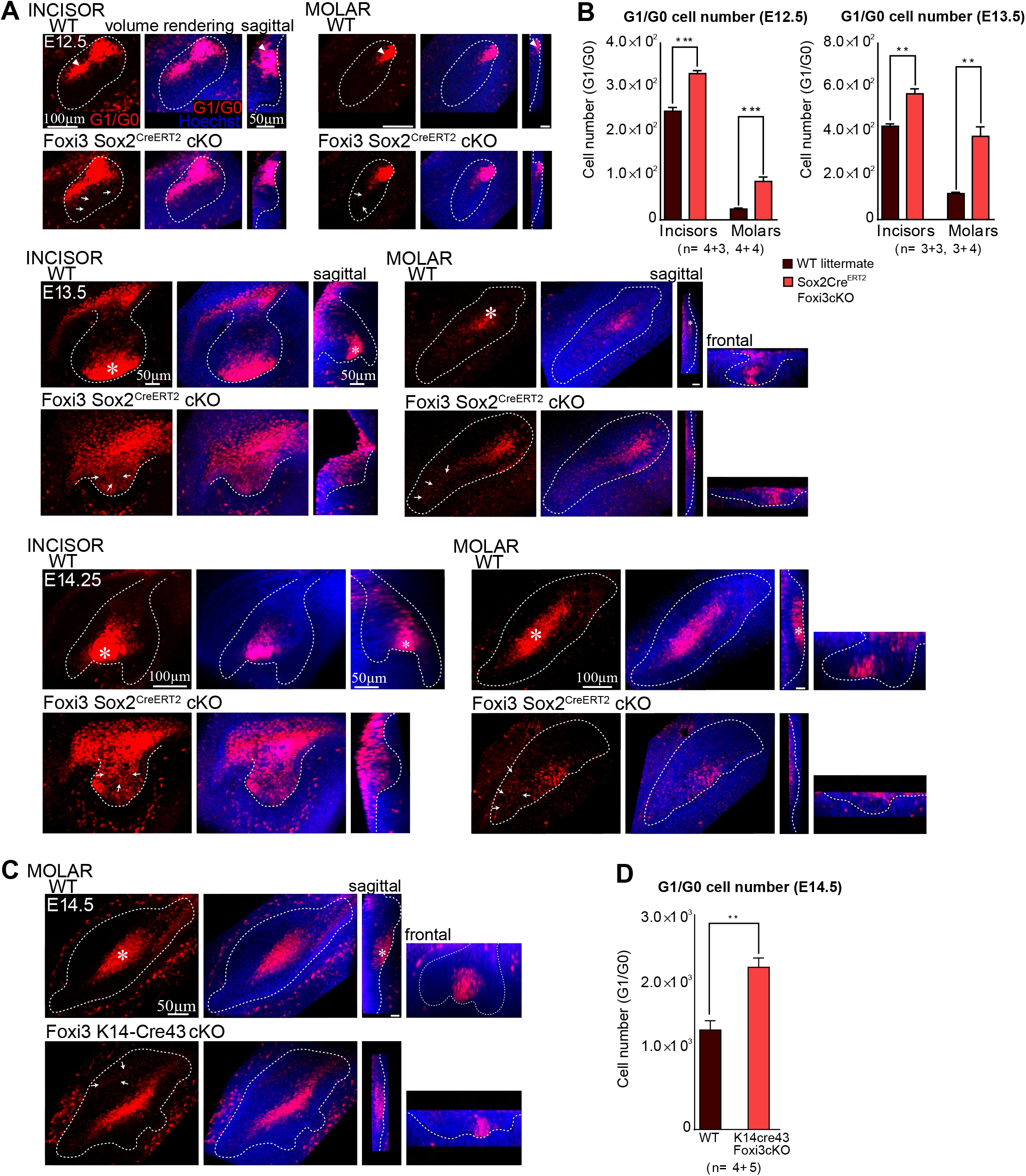
Abnormal cell cycle exit and growth impairment in the Foxi3cKO mutants. The Fucci G_1_/G_0_ reporter can be used to identify signaling centers in early developing incisors and molars. (A-B) Whole-mount confocal fluorescence images of the cell cycle indicator (Fucci) for G_1_/G_0_ phase cells (red) of WT and mutant teeth. Nuclei were stained with Hoechst 33342 (blue). Volume renderings from planar view, and sagittal and frontal optical sections. The frontal sections in z plane are arranged perpendicularly from the sagittal sections rotating around the pEK as axis. Epithelial borders are delineated with a dotted line. IK (arrowhead) and emerging EK (asterisk). In WT teeth, confined G_1_/G_0_ phase cells were present in the signaling center regions (IK and EK) in incisors and molars of stages E12.5, E13.5, E14.25 and E14.5. More specifically, IK G_1_/G_0_ Fucci cells comprised the IK in WT and mutant teeth at the labial side, in the frontal part of incisor and molar buds. The EK G_1_/G_0_ Fucci cells located at the deepest part of the WT incisor and molar buds, whereas in mutant teeth there was no evident EK structure. (A) In all stages studied the Sox2^CreERT2^Foxi3cKO mutant teeth show more widely distributed G_1_/G_0_ phase cells scattered throughout the IK and EK regions. The normally proliferating bud populations showed abnormal cell cycle exit (arrows). (B) Quantification of cell number of Fucci G_1_/G_0_ positive cells in WT and mutant teeth of stages E12.5 and E13.5 teeth (N_E12.5_=4+3, 4+4; N_E13.5_=3+3, 3+4). In both stages, mutant incisors and molars display a higher number of cells in G_1_/G_0_. Altogether, mutant tooth growth was arrested in late bud stage (E14.5) probably due to the high number of cells in non-proliferative state. (C) Cell cycle exit dynamics was also studied in the bud stage K14^Cre43^ Foxi3 conditional knockout mutant E14.5 molars. In E14.5, WT molars show a new G_1_/G_0_ focus corresponding to the EKs, whereas the G_1_/G_0_ population in the Foxi3cKO molars is more widespread all over the bud. (D) Quantification of cell number of Fucci G_1_/G_0_ positive cells in WT and K14^Cre43^Foxi3cKO mutant teeth (N_E14.5_=4+ 5). At E14.5, mutant molars display a higher number of cells in G_1_/G_0_. For all statistical tests: mean±sem, nonparametric Mann-Whitney *U*-test, p<0,05 *, p<0,01**, p≤0.001***.

We next looked at the non-proliferative population of E14.5 molars in the bud stage K14^Cre43^ Foxi3 conditional knockout mutant model. In WT teeth, the pEK signaling center was located at the center of the molar deep in the bud whereas in K14^Cre43^Foxi3cKO molars the G_1_/G_0_ cell population (presumably pEK cells) were more widely distributed (Figure 3c). Quantifications of Fucci G_1_/G_0_ cell number resulted in a higher amount of non-proliferating cells in E14.5 mutant molars compared to WTs (Figure 3d). Thus, the extra epithelial foldings observed in E14.5 mutant molars are likely to be caused by the abnormal localization and increase of G_1_/G_0_ cells.

### Decreased apoptosis in signaling centers in Foxi3cKO mutants

Silencing through apoptosis is a hallmark of signaling centers in different organs (Matalova et al., 2004; Nonomura et al., 2013, Vaahtokari et al., 1996). Apoptotic activity has also been noted in the incisor and molar IKs (Munne et al., 2009; Vaahtokari et al., 1996; Mogollón et al., 2021). We next studied apoptosis of signaling centers in the early stage Sox2^CreERT2^ Foxi3 conditional knockout mutants. We chose the stages between E12.75-E13.5 when apoptotic activity in both IK and EK signaling centers has been reported. Apoptotic cells were detected with immunofluorescence staining of cleaved caspase 3 (Casp3).

At E12.75, WT teeth showed apoptotic cells mainly restricted to the IK area whereas in mutant teeth almost no apoptosis was seen in the IK region (Figure 4a). Already at E13.5, apoptosis was more prominent in the IK from WT teeth than one day before and in Sox2^CreERT2^Foxi3cKO mutant teeth it remained scarce (Figure 4b). Quantification of apoptosis, measured as % of apoptotic Cas3+ cells of Fucci G_1_/G_0_+ cells, showed that in the stages studied (between E12.75-E13.5), apoptosis was reduced in both mutant incisors and molars (Figure 4c). Quantifications of the absolute number of apoptotic cells also resulted in a reduced number in mutant teeth (data not shown). Interestingly, incisors and molars showed different apoptosis patterns: both WT and mutant incisors displayed a % of apoptotic cells at E12.75 that further increased at E13.5. Whereas in molars, E12.75 WT teeth showed most G_1_/G_0_ cells going through apoptosis, and this % decreased by E13.5; on the other hand, apoptosis did not change significantly in Sox2^CreERT2^ mutant molars (Figure 4c). We conclude from this evidence that the early conditional knockdown of Foxi3 from E11 onwards in dental epithelia affects signaling center silencing and overall tissue homeostasis.

**Figure 4.**
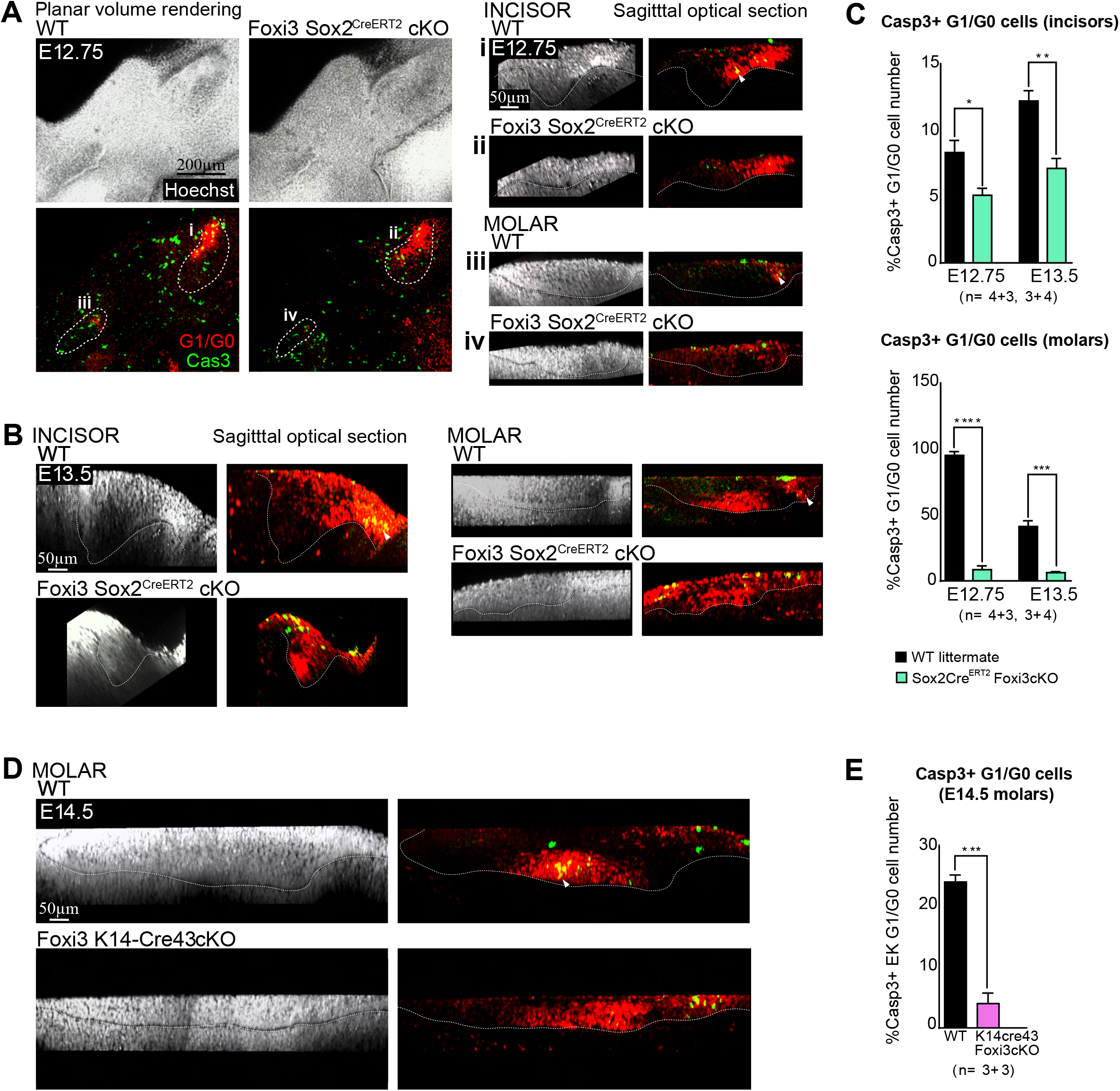
Decreased apoptosis in signaling centers in the Foxi3cKO mutants. For detection of apoptotically active cells the tissues were immunofluorescence stained with an antibody specific for cleaved caspase 3. Whole-mount confocal fluorescence images of the cell cycle indicator (Fucci) for G_1_/G_0_ phase cells (red) and cleaved caspase 3 immunofluorescence staining (Cas3, green). Nuclei were stained with Hoechst 33342 (white). Epithelial bud (dotted line) and Cas3+ cells (arrowhead). Volume renderings from planar view and optical sections from sagittal side view of WT and mutant teeth. G_1_/G_0_ phase cells were present in the signaling center regions (IK and EK) in WT incisors and molars at (A) E12.75 and (B) E13.5, whereas in Sox2^CreERT2^ Foxi3cKO mutant incisors and molars the G_1_/G_0_ phase cells were more widely distributed and scattered throughout the IK and EK regions and bud populations. Apoptotic cells were mainly confined to the signaling centers (IK and EK) in WT teeth, whereas in mutant teeth apoptosis was reduced. (C) Quantification of percentage of G_1_/G_0_ cells going through apoptosis (Cas3+ cells) in WTs and Sox2^CreERT2^ mutants E12.75 and E13.5 teeth (N_incisors_=4+3, 3+4; N_molars_=4+3, 3+4). This quantification shows that a lower percentage of Fucci G_1_/G_0_ positive cells is going through apoptosis in Sox2^CreERT2^ mutant teeth. (D) Apoptosis was also visualized in the bud stage K14^Cre43^Foxi3 conditional knockout mutant molars at E14.5. Apoptotic cells were mainly confined to the pEK signaling center in WT molars, whereas in K14^Cre43^Foxi3cKO mutants’ apoptosis was reduced. (E) Quantification of percentage of G_1_/G_0_ cells going through apoptosis (Cas3+ cells) in WT and K14^Cre43^Foxi3cKO mutant E14.5 molars (N_E14.5_=3+3). This quantification shows that a lower percentage of Fucci G_1_/G_0_ positive cells is going through apoptosis in K14 ^Cre43^Foxi3cKO mutant molars. For all statistical tests: mean±sem, nonparametric Mann-Whitney *U*-test, p<0,05 *, p<0,01**, p≤0.001***. All in all, a lower percentage of Fucci G_1_/G_0_ positive cells is going through apoptosis mutant teeth in both Foxi3cKO models between E12.5-E14.5.

We studied apoptosis of signaling center cells in the bud stage K14^Cre43^ Foxi3 conditional knockout mutants (Figure 4d). We studied molars at E14.5 when there is a clear difference in phenotype of the bud stage K14^Cre43^ Foxi3 conditional knockout mutants compared to WT molars (Jussila et al. 2015). Casp3 staining in Fucci G_1_/G_0_ WT teeth showed apoptotic cells mainly restricted to the pEK area, whereas in mutant teeth almost no apoptosis was seen in that region (Figure 4d). Quantification of % of apoptotic Cas3+ cells (of Fucci G_1_/G_0_+ cells) showed that apoptosis was reduced in E14.5 K14^Cre43^Foxi3cKO mutant molars compared with WT molars (Figure 4e). Hence, the apoptosis decrease was observed in both Foxi3cKO mutant models.

### Abnormal proliferation patterns in Foxi3cKO mutant teeth cause epithelial size and shape differences

Epithelial tooth budding in WT incisors and molars occurs via cell proliferation regulated by non-proliferative signaling centers (Ahtiainen et al., 2016; Mogollón et al. 2021). Since we identified an increase in cell cycle exit, we wanted to know the effect of Foxi3 knock down in cell proliferation. Cell proliferation was studied with Fucci bitransgenic mice which express nuclear red in G_1_/G_0_ phase (Cdt1-mKO) and nuclear green in S/G_2_/M phases (Gem-mAZ) (Sakaue-Sawano et al., 2008). We studied the effects of cell proliferation dynamics on epithelial tooth bud invagination in WT and both Foxi3cKO mutant models at placode (E12.5), bud (E13.5) and early cap stage (E14.5) (Figure 5). 3D epithelial renderings of the epithelial placode/bud were done by manually drawing the epithelial-mesenchymal boundary visualized with Hoechst and contouring the epithelium. The epithelial renderings defined the location and distribution of epithelial cells in G_1_/G_0_ and S/G_2_/M phases. To make 3D surface renderings of single nuclei, nuclei expressing either Fucci reporter of G_1_/G_0_ or S/G_2_/M cell cycle phases were used.

**Figure 5.**
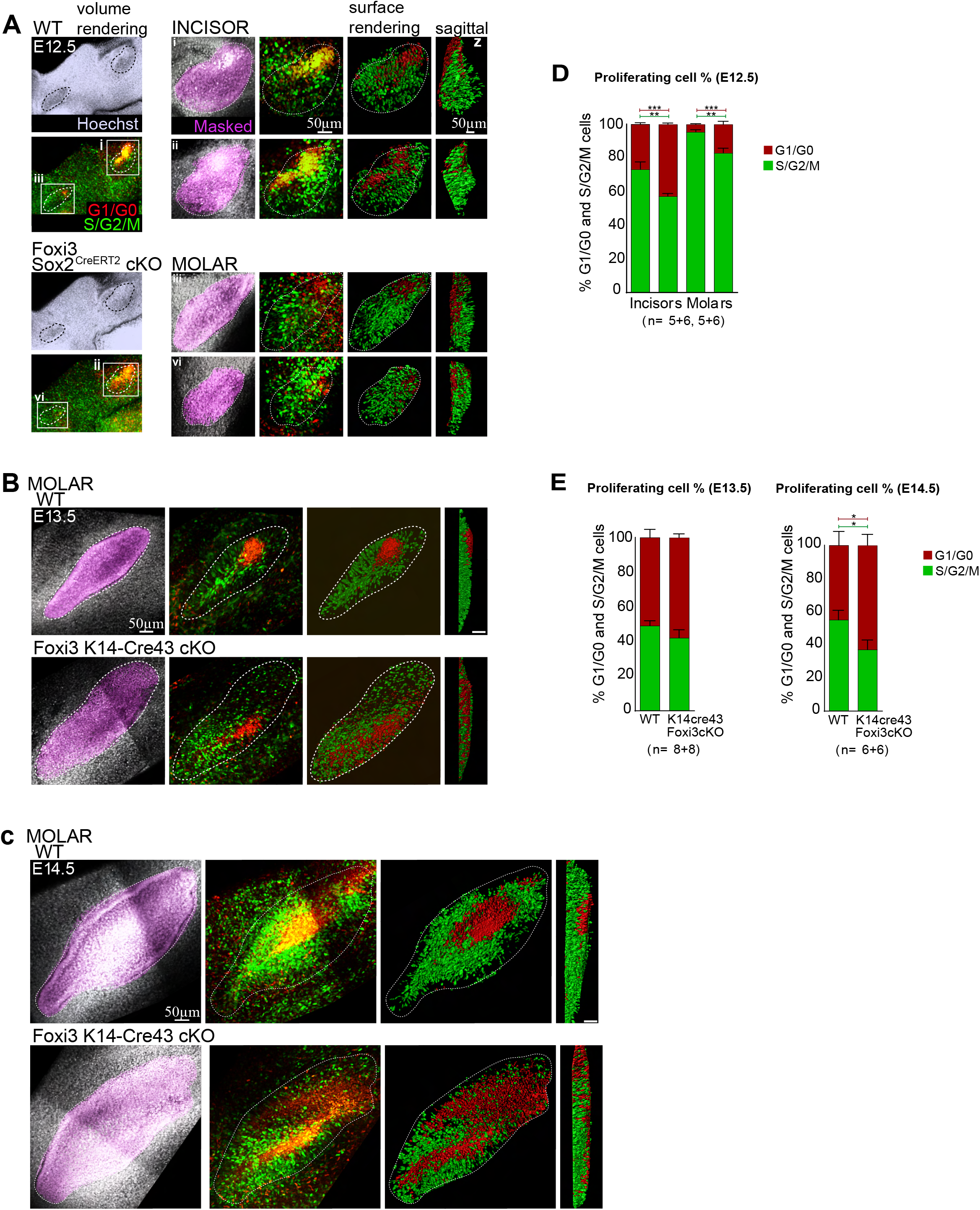
Proliferation pattern is abnormal in Foxi3 conditional mutants which causes epithelial size and shape differences. Fucci reporters are used to monitor cell cycle status. Whole-mount confocal fluorescence images of the Fucci reporters for G_1_/G_0_ (red) and S/G_2_/M (green) phase cells at different stages of the Foxi3 mutants and WT teeth. Nuclei stained with Hoechst 33342 (white). Epithelial bud (dotted line). Volume renderings from planar view and optical sections from sagittal view. Planar volume renderings of masked dental epithelia in respect to body axes (magenta). In green and red, surface renderings of the nuclei in G_1_/G_0_ and S/G_2_/M cell cycle phases in the developing epithelial bud of incisors and molars (planar and sagittal views). G_1_/G_0_ phase cells were present in the signaling center regions (IK and EK) in WT incisors and molars at stages (A) E12.5, (B) E13.5, and (C) E14.5. On the other hand, in E12.5 Sox2^CreERT2^ Foxi3cKO incisors and molars (A), and E13.5-E14.5 K14^Cre43^ Foxi3cKO molars (B and C), the G_1_/G_0_ phase cells were more widely distributed and scattered throughout the buds. The S/G_2_/M cell populations in WT teeth localized in both basal and suprabasal compartments, mainly closer to the signaling centers, whereas in mutant teeth the proliferative cell populations were also more widely distributed throughout the buds. (D) Quantification of the percentage of epithelial cells in G_1_/G_0_ and S/G_2_/M stages in E12.5 WT and Sox2^CreERT2^ Foxi3cKO mutant incisors and molars (N_E12.5_=5+6, 5+6). Mutant teeth show a lower percentage of proliferative cells in both incisors and molars compared to the WT counterparts, and a higher percentage of cells in non-proliferative state. (E) Quantification of the percentage of epithelial cells in G_1_/G_0_ and S/G_2_/M stages in E13.5 and E14.5 WT and K14^Cre43^ Foxi3cKO mutant molars (N_E13.5_=8+8, N_E14.5_=6+6). Mutant teeth show a lower percentage of proliferative cells compared to the WT counterparts; and a higher percentage of cells in non-proliferative state Notably, percentage of cells in proliferative phase increase as tooth epithelium grows in WT teeth between E13.5-E14.5, whereas it decreases in mutant tissue. For all statistical tests: mean±sem, nonparametric Mann-Whitney *U*-test, p<0,05 *, p<0,01**, p≤0.001***.

In all stages analyzed, the proliferative cell populations in WT teeth were localized both in basal and suprabasal compartments, mainly closer to the signaling centers (the G_1_/G_0_ cell populations; Figure 5a-c). In contrast, E12.5 Sox2^CreERT2^Foxi3cKO mutant teeth showed more scattered S/G_2_/M cell populations around the presumable IK regions, as well as the G_1_/G_0_ cell populations when compared to WT E12.5 teeth (Figure 5a). Similarly, the K14^Cre43^Foxi3cKO mutant model displayed E13.5 and E14.5 molars with more scattered S/G_2_/M cells around the region where the pEK would normally reside at the center/bottom of the bud (which also comprised a wider G_1_/G_0_ cell distribution), when compared to WT E13.5-E14.5 molars (Figure 5b-c). Quantification of the proportions of cells in different cell cycle phases (measured as % of cells in G_1_/G_0_ phase and % of cells in S/G_2_/M phase) at E12.5, E13.5, and E14.5 stages indicated a higher portion of Fucci G_1_/G_0_ cells in mutant incisors and molars than in WT tissue, in which more cells were in the proliferative S/G_2_/M phase compared to mutant tissue (Figure 5d-e). All in all, mutant teeth display a widespread distribution of S/G_2_/M cells throughout the bud epithelia, as in the case of the more widely distributed G_1_/G_0_ cells (Figure 5a-c). Likewise, the observed differences in proliferation and cell distribution and number could account for the differences in tooth epithelium shape and size. Overall, Foxi3cKO mutant teeth are smaller, flatter, and abnormally shaped likely caused by proliferation dynamics differences.

### Cell fate changes of signaling center and bud cells in Foxi3cKO mutants

To further verify the signaling center identity of the Fucci G_1_/G_0_ cells broadly distributed in mutant teeth, we used the fluorescent canonical Wnt signaling reporter TCF/Lef:H2B-GFP in E13.25-E14.5 mandibles (Figure 6).

**Figure 6.**
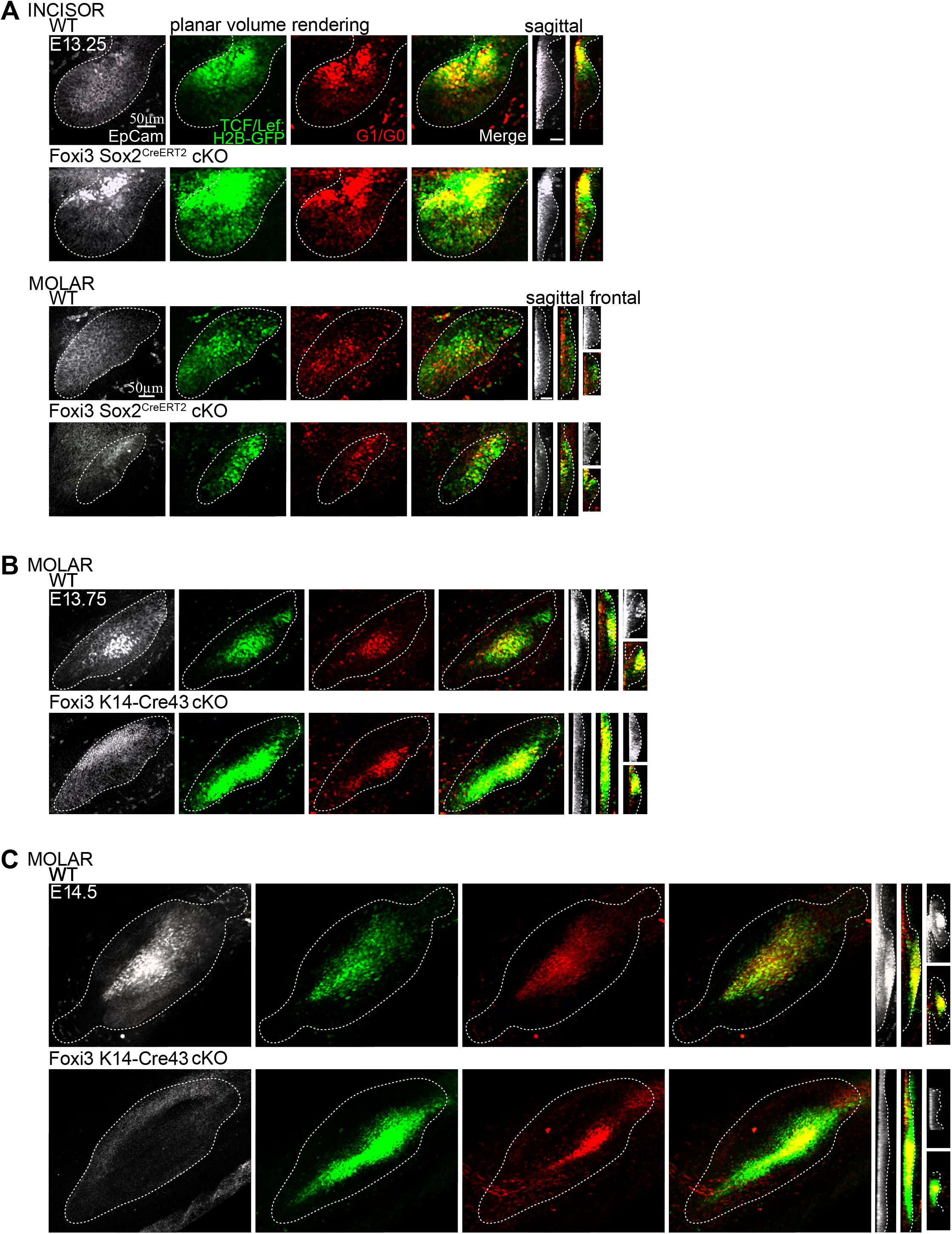
Cell fate changes in Foxi3cKO mutants (increased WntHi cells surrounding the G_1_/G_0_ cell populations) The canonical Wnt signaling reporter TCF/Lef:H2B-GFP expressing WntHi cells is closely juxtaposed to G_1_/G_0_ signaling center cells, as two different cell populations in close contact with each other through normal tooth bud development. Whole-mount confocal fluorescence images of the cell cycle indicator (Fucci) for G_1_/G_0_ phase cells (red) and TCF/Lef:H2B-GFP (green). Immunofluorescence staining of the epithelium (EpCam, grey, dotted line). (Planar view) Each figure is a volume rendering of the xy plane. (Sagittal and frontal view). Each figure is an optical section in z plane of WT and mutant incisors and molars. (A) At E13.25, WntHi cells were present in the remaining IK cells in WT incisors in a more restricted manner compared to Sox2^CreERT2^Foxi3cKO mutant incisors. Mutant incisors displayed a more widespread distribution of both Fucci G_1_/G_0_ and TCF/Lef:H2B-GFP cells in the IK and all over the buds. In WT molars, the emerging pEK with G_1_/G_0_ cells were surrounded by WntHi cells, distributed centrally. In Sox2^CreERT2^Foxi3cKO mutant teeth, G_1_/G_0_ and WntHi cell distribution is more spread and scattered all over the bud. K14^Cre43^Foxi3cKO mutant (B) E13.75 and (C) E14.5 molars also displayed a more widespread distribution of both Fucci G_1_/G_0_ and TCF/Lef:H2B-GFP cells all over the bud.

Our results revealed that E13.25 Sox2^CreERT2^Foxi3cKO mutant teeth showed more intense GFP+ signals around the presumable IK and EK regions, as well as the G_1_/G_0_ cell populations when compared to WT E13.25 teeth (Figure 6a). Similarly, the K14^Cre43^Foxi3cKO mutant model displayed E13.75 and E14.5 molars with more intense arrangement of WntHi cells around the presumable pEK when compared to WT E13.75-E14.5 molars (Figure 6b-c). In summary, the TCF/Lef:H2B-GFP reporter (i.e., WntHi cells) was detected in proximity with the G_1_/G_0_ cell population located in the signaling center regions. Mutant teeth displayed an intense, widespread distribution of WntHi cells throughout the buds, likely because G_1_/G_0_ cells were also more widely distributed within the tooth epithelia (Figure 6a-c). Altogether, these data verified that the expanded Fucci G_1_/G_0_ cell populations seen in mutant teeth have signaling center identity, and that mutant G_1_/G_0_ signaling centers are thus expanded in distribution and in signaling molecules. Independently of the stage and mutant model, we could conclude that Foxi3 deficiency results in an increase and abnormal distribution of signaling center cells, and at the same time the normally proliferating bud cells switch to signaling center fate.

Sox2 is a marker for pluripotency during development. In teeth, it is firstly expressed throughout the entire dental lamina and then becoming progressively restricted to the incisor and molar regions. At bud stage, Sox2-positive cells are restricted to the bud proliferative populations, mainly confined to the lingual side and not within the signaling centers (Juuri et al., 2013; Maria Sanz-Navarro et al., 2018; Ahtiainen et al., 2016). We lastly wanted to elucidate if the mutant bud cells, which have abnormally shifted to signaling center fate and exited the cell cycle, also stopped expressing Sox2. Whole-mount immunostainings of Sox2 on WT and K14^Cre43^Foxi3 E14.5 molars showed that cells that exhibited cell cycle exit (as seen with the Fucci G_1_/G_0_ reporter) and expressed signaling center markers, also did not show Sox2 expression (Figure S2).

## Discussion

Early tooth morphogenesis requires an epithelial initiation knot (IK) signaling center for the progression of both incisor and molar tooth development from placode to bud stages (Ahtiainen et al., 2016; Mogollón et al., 2021). However, little has been studied regarding what happens when this IK signaling center is misregulated. Since tissue recombination studies have indicated that the epithelium carries the initial capacity for instructing tooth morphogenesis (Lumsden, 1988; Mina and Kollar, 1987), generating a model system that allows genetic manipulation of the early dental epithelia is important for the understanding of tooth initiation and IK regulation. In the present study, we took advantage of the inducible Cre model system Sox2^CreERT2^ and explored early development misregulation by knocking down the early dental epithelial marker Foxi3. Here we show that (1) the Sox2^CreERT2^ model is a suitable tool for studying early dental morphogenesis; (2) the Foxi3 transcription factor is essential for early tooth morphogenesis; and (3) by studying Foxi3 we can investigate signaling center function.

Sox2 and Foxi3 show similar expression patterns in the epithelial dental lamina, coinciding at E11.5 with the initiating IK signaling centers (Ahtiainen et al., 2016). Later in the forming buds, Foxi3 and Sox2 expression patterns become more restricted to the lingual side within the bud population (Juuri et al., 2013; Maria Sanz-Navarro et al., 2018; Ahtiainen et al., 2016), indicating that the more mature IK cells no longer express these markers. Such data reveals that the IK cells have become differentiated, knowing that Sox2 and Foxi3 are markers of epithelial cells in undifferentiated/progenitor state as indicated in previous studies (Juuri et al., 2013; Jussila et al. 2015). The wide-ranging expression pattern of Foxi3 along several stages of tooth morphogenesis and diverse cell populations suggests that this molecule might have multiple roles depending on the context (Shirokova et al., 2013). Therefore, it was expected that the phenotype displayed by the bud stage K14^Cre43^ Foxi3 conditional knockout mutant model developed by Jussila et al. (2015) would differ from the one caused by the early Sox2^CreERT2^ Foxi3 conditional knockout mutant model. Some similarities between the two models include the increase of the G_1_/G_0_ cell population upon Foxi3 deficiency (observed in the previous study as p21 upregulation), increase in signaling center marker expression, and decreased epithelial downgrowth.

On the other hand, the differences in timing of the Foxi3 knockdown in the two mutant models accounted for some major differences. The early stage Sox2^CreERT2^ mutant model hampers Foxi3 expression at the time it is a marker of the IK at E11.5. By the time it should switch to the bud proliferative population and define the different cell identities at E12.5 (IK = differentiated = no Foxi3 vs. bud proliferative cells = undifferentiated = Foxi3 present) its deficient expression alters the balance in cell populations causing most cells to attain signaling center identity. Therefore, if too many cells acquire non-proliferative signaling center fate, the growth of dental buds gets arrested along with further tooth development in both incisors and molars (Figure 7). In the case of the bud stage K14^Cre43^ mutant model, Foxi3 expression is efficiently knocked down around E13 when the Foxi3 is already switching to be a marker for the bud proliferative cell population. This specific timing turns some of the bud cells to switch to signaling center identity but not all of them and causes an expanded signaling center area. The abnormal distribution of signals results in scattered proliferation patterns that further affect the overall molar tooth shape (Figure 7). The atypical morphology of the presumptive pEK could ultimately disrupt the distribution of sEKs further in development, as shown by Jussila et al. (2015). Early processes may underlie the adult molar phenotype of the K14^Cre43^ mutants: fused and flatter molars with altered and shallower cusps (Jussila et al., 2015). Similarly, the decrease in Sox2 expression also suggested that some K14^Cre43^ Foxi3cKO mutant molar cells may now lack pluripotency due to deficiency of Foxi3 (Figure S2).

**Figure 7.**
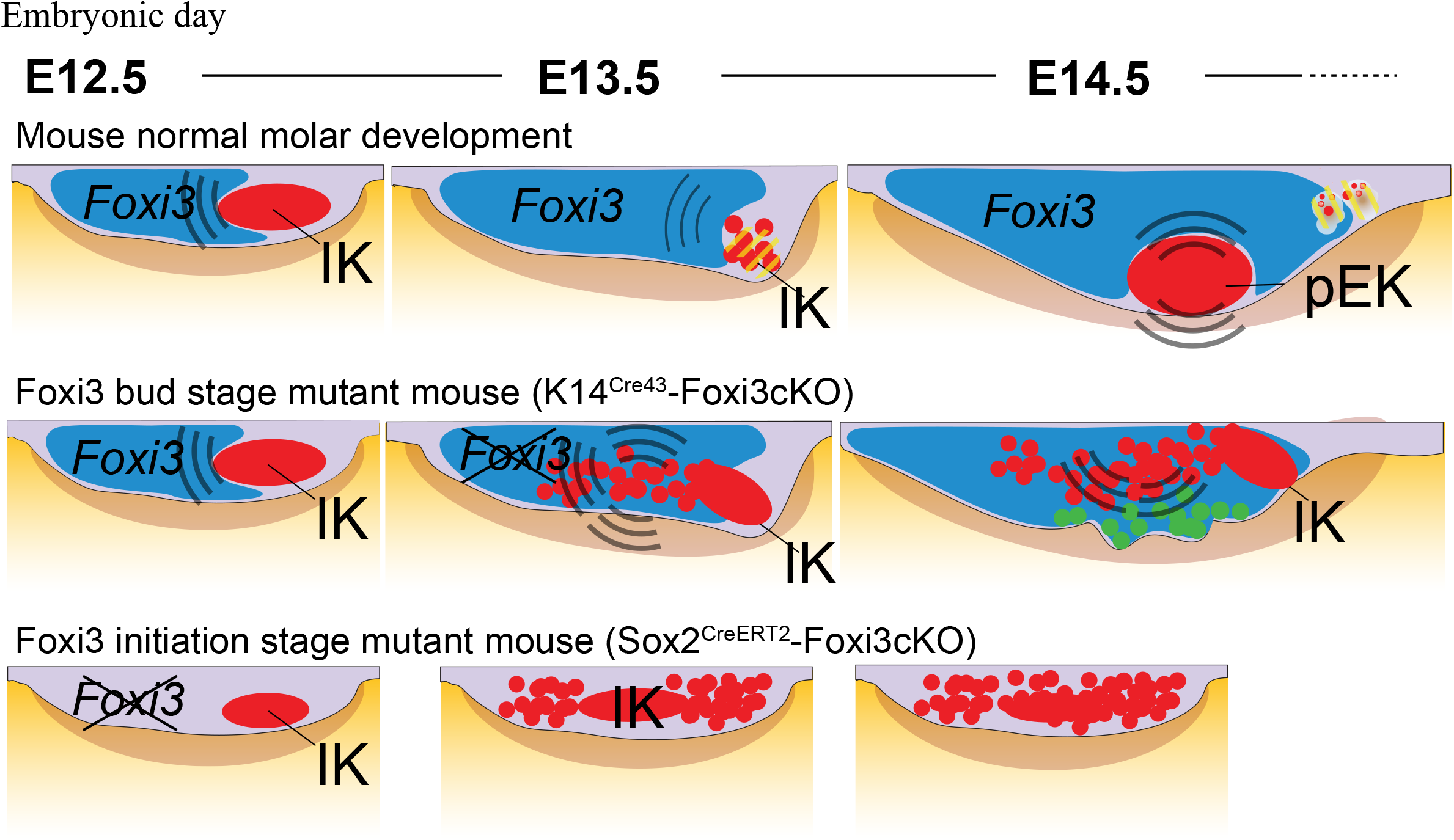
Cell mechanisms underlying Foxi3 knock down in Sox2^CreERT2^ and K14^Cre43^ mutant models. Schematic of the cellular dysplasia mechanisms occurring in early tooth development due to Foxi3 knock down. (A) In normal development Foxi3 is expressed in the whole placode and later is restricted in the bud cell population, excluding the IK signaling center cells. (B) Upon conditional deletion of Foxi3 at placode stage (E11), the proliferative bud population changes in cell fate definition and most cells switch to signaling center identity, causing increased cell cycle exit, prolonged maintenance of signaling center signals, and impaired growth. (C) Upon conditional deletion of Foxi3 at bud stage (E13), fewer number bud cells and a more mature population of these cells switch to signaling center fate and teeth can develop, however, resulting in abnormal bud shape and ultimately fused molars. In summary, changes in signaling center fate definition, cell cycle changes and prolonged signaling center maintenance, suggests a novel cellular disease mechanism in ectodermal dysplasia.

The fact that in this model the molars are the most clearly affected indicates that the timing of Foxi3 knockdown is crucial in the different tooth types around stage E13, when Foxi3 starts to be predominantly a marker of the bud cell population. The less affected incisor development in the bud stage K14^Cre43^ model could be caused by distinctions in signaling center dynamics and bud proliferation rate differences. Altogether, Foxi3 deficiency in early tooth formation results in a switch in cell identity towards the more differentiated, signaling center fate. Likewise, the Sox2^CreERT2^ system showed to be a novel and suitable model for genetic and functional manipulation of the IK signaling center in both murine incisors and molars during the early stages of tooth morphogenesis. Conversely, the K14^Cre43^ system is a useful tool to study changes in EK dynamics, bud-to-cap morphogenesis and beyond.

Mogollón et al. (2021) and our early Sox2^CreERT2^ Foxi3 mutant model evidenced that the function of the IK is to drive enough bud proliferation to build a proper epithelial bud for tooth morphogenesis to proceed normally. If the IK is either underdeveloped or abnormally overdeveloped, this leads to a scarce number of proliferative bud cells causing a growth arrest at bud stage. Jussila et al. (2015) already suggested that Foxi3 is involved in cell identity specification. Moreover, Foxi3 mutant phenotypes in our Sox2^CreERT2^ model showed a continuous pattern (presumably caused by variability in tamoxifen efficiency): a phenotypic spectrum from very severe (arrested small bud), intermediate (bigger arrested bud), to less severe (small cap). Jussila et al. (2015) also reported that variability in Foxi3 levels throughout evolution could be a mechanism of cusp pattern modification, as it has been reported for other tooth genes with comparable phenotypic continuity (Pispa et al. 2004; Harjunmaa et al. 2012). Our analyses of the K14^Cre43^Foxi3cKO model may suggest that the main role of the murine EKs/pEKs is to regulate tooth shape, and alterations in the EK result in abnormal formation of sEKs, cusp pattern defects and fused molars. Therefore, from these findings we hypothesize that the two different signaling centers (IK and EK) have similar behaviors and identity, and they both induce proliferation to the neighboring bud cell populations; yet we speculate they produce different morphogenetic effects which are context dependent. Namely, we suggest that the main role of the IK is to regulate tooth size, and the EK tooth shapes. Thus, the processes controlling size are separated from those involved with shape. This was observed with the fact that in the Sox2^CreERT2^ model there was a more severe reduction in size but not entirely changes in shape, whereas with the K14^Cre43^ model there was a much more prominent phenotype change in tooth shape, compared to tooth size. Li et al. (2016) studied the mechanisms involved in early molar morphogenesis and discovered that the process of stratification (cell layer formation) and invagination (budding) are separate processes: inhibition experiments of Fgf and Shh pathways resulted in either differently shaped (flatter) or smaller bud. Dental complexity studies have previously proposed that increase in cusp number is independent of tooth size during evolution (Evans et al., 2007; Harjunmaa et al., 2012).

The increase in number of signaling center cells in the Foxi3cKO models was accompanied by an increment in signaling center signal (e.g., canonical Wnt,), a decrease in progenitor cell markers (Sox2), and a decrease in apoptosis (seen by reduced cleaved caspase-3 expression). We suggested in our previous study that it is possible that silencing of signaling centers through apoptosis is caused by the downregulation of canonical Wnt and Shh signals (Mogollón et al., 2021), so it is plausible that the abnormal prevalence of these signals upon Foxi3 deficiency in the mutants hampers the proper silencing of signaling centers (Jussila et al., 2015). Previous research indicates that Fgfs and Shh can also have protective roles against apoptosis (Vaahtokari et al., 1996; Cobourne et al., 2001). The caspase-3^-/-^ mouse model resulted in a molar (at cap stage) with a resembling phenotype as the Foxi3cKO mutants: disorganized epithelium across the pEK area and abnormal expression of signaling center signals (Matalova et al., 2003). Along with cell quiescence and silencing through apoptosis, cell migration and condensation are hallmarks of ectodermal signaling centers. IK signaling center condensation via cell migration is essential for tooth budding, and this condensation is regulated by the presence of Wnt10b signals and cells with TCF/lef:H2B-GFP (WntHi+) identity (Ahtiainen et al., 2016; Mogollón et al., 2021). The observed expansion of the canonical Wnt reporter TCF/lef:H2B-GFP and Fucci G_0_/G_1_ reporter domains in the Foxi3 mutants could be partially related to defective condensation. Li et al. (2016) proposed that Shh drives cell rearrangements and movements. Jussila et al. (2015) concluded that Foxi3 is involved in the formation of the suprabasal cell population, since K14^Cre43^ mutant molars buds look shallower with reduced number of these cells. In the present study, we have examined that one of the mechanisms causing such a phenotype is the lack of proliferation due to increased cell cycle exit; however, we do not rule out the possibility that an impairment in cell rearrangements could also cause the lack of normal suprabasal cell formation. Moreover, although speculative, it is possible that if signaling center apoptosis is induced by contact inhibition, the lack of mechanical crowding of the signaling center cells in mutant teeth could also account for the decreased apoptosis observed. Further studies, with live imaging approaches, would shed light concerning these processes.

Foxi3 function can vary greatly depending on the context, and it has shown to have potential roles during evolution (Iurlaro et al., 2013; Jussila et al., 2015). The possible evolutionary function of Foxi3 is supported by the fact that its expression ranges across different organs and phylogenetic classes: it is frequently expressed in the early stages of organ morphogenesis, for instance, in murine mammary gland and hair follicle placodes, as well as in cranial placodes from chick embryos (Shirokova et al., 2013; Khatri et al. 2014). Notably, as in our mutant teeth, loss of Foxi3 in hair follicles results in impaired downgrowth, cell specification and downregulation of stem cell identity (Shirokova et al., 2016). It has been shown that Foxi3 is a downstream target of Eda, suggesting that the phenotype seen in dogs with heterozygous mutation of Foxi3 resembles the human ectodermal dysplasia (HED), displaying abnormal hair and teeth (Shirokova et al., 2013). However, from our data it is now evident that even though Foxi3 can be regulated by the Eda pathway, the phenotypes observed when Eda is knockdown vs. Foxi3 are distinctive. Whereas loss of Eda results in reduced signaling center cell number (Ahtiainen et al., 2016), Foxi3 knockdown results in an increased number of cells with signaling center cell identity (however not enough proliferative cells available). In that case, the phenotype caused by early knockdown of Foxi3 would resemble more the phenotype observed when Eda pathway is hyperactivated. More specifically, mutant mice overexpressing different components from the Eda pathway display tooth shape and number defects (Mustonen et al., 2003; Pispa et al., 2004). For instance, ectopic EdaR expression outside the signaling centers (using the K14 promoter) impairs tooth patterning, cusp number, and differentiation of ameloblasts and enamel formation, depending on the extent of transgene expression (Pispa et al., 2004). Hence, in the context of early tooth development Foxi3 might also be regulated by other pathways; for example, Activin A, which can upregulate Foxi3 mRNA levels *in vitro* (Shirokova et al., 2013).

Similar to the findings shown by Jussila et al. (2015) reporting that Foxi3 regulates Wnt signaling, changes in the signaling centers observed in our Foxi3cKO mutants resemble the ones reported for *Catenb*^Δex3k14/+^ mutant mice with hyperactive Wnt signaling (Järvinen et al., 2006): extra epithelial foldings, increase of p21 and Shh, and apoptosis decrease. Nonetheless, these mice display extra teeth, possibly due to higher amount of proliferative bud cells. In parallel, in knockout mice lacking the downstream molecule of canonical Wnt signaling Lef1, the tooth germ is arrested at bud stage like in the Sox2^CreERT2^ mutants (van Genderen et al., 1994). All in all, a very precise equilibrium of signaling molecules determines the induction or inhibition of tooth morphogenesis and is essential for proper formation of shape, size and number.

In conclusion, we developed a new model for the genetic manipulation of tooth development at placode stage, with the Sox2^CreERT2^ transgenic mouse, opening the possibility for further research that has been limited in the past. We suggest that the Foxi3 transcription factor is required for cell identity determination between the IK signaling center and the bud proliferative cell population during early incisor and molar tooth morphogenesis. The results of this study corroborate the importance of properly functioning IK and EK signaling centers for tooth downgrowth and shape, requiring a specific size, positioning and silencing in order to generate normal murine tooth bud.

## Supplemental Figure Legends

**Figure S1.**
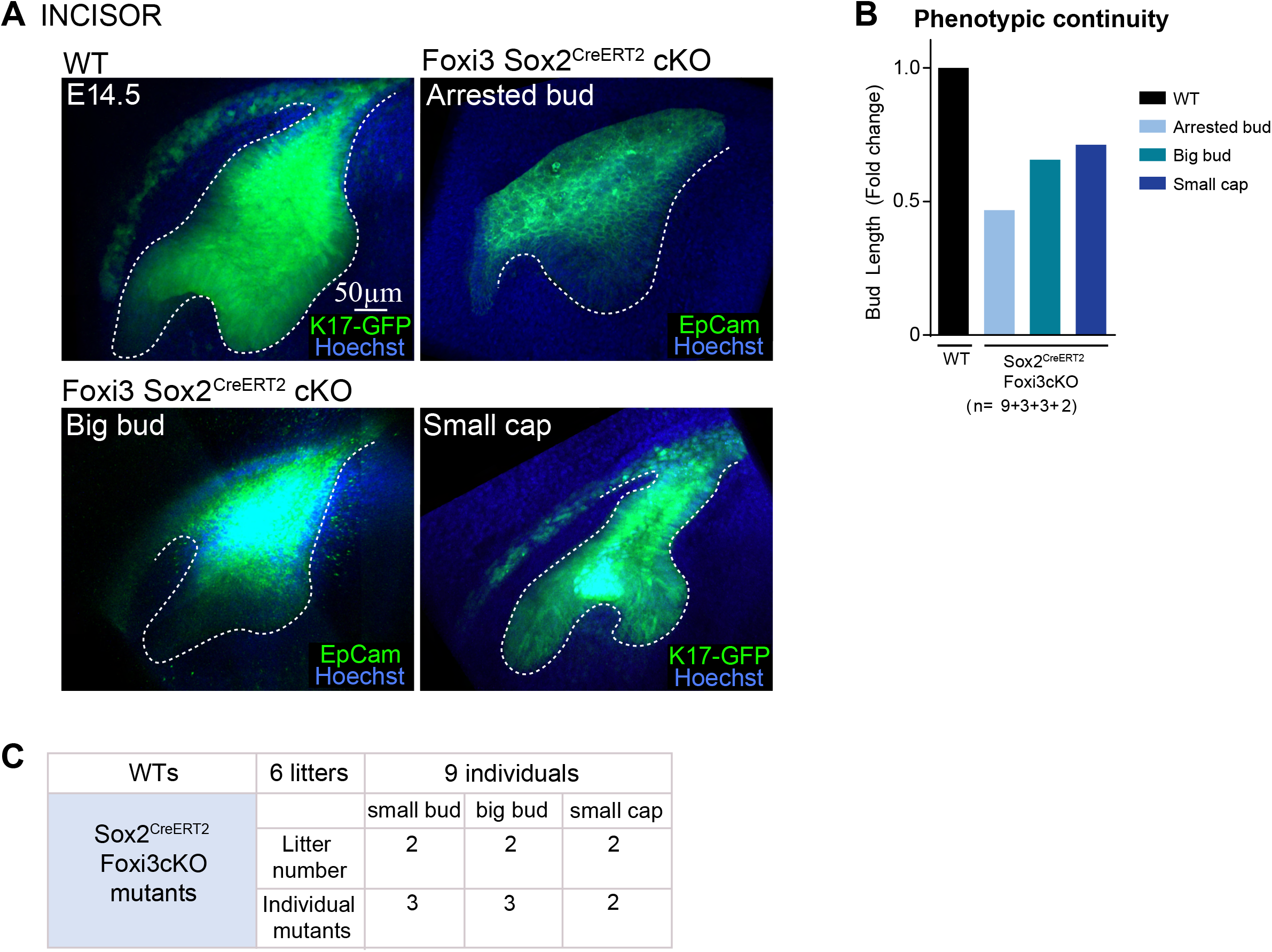
Phenotypic continuity in Sox2CreERT2 mutants at arrested bud-to-cap transition. Depending on the efficiency of tamoxifen induction in the Sox2^CreERT2^Foxi3cKO model, we observed a range of phenotypes caused by varying levels of Foxi3 knock down. Whole-mount confocal fluorescence images incisors and molar epithelial buds with keratin 17-GFP (green) or EpCam (green). (Planar view) Each figure is a volume rendering of the xy plane. Tooth epithelium perimeter (dotted line). Nuclei were stained with Hoechst 33342 (blue). (A) Sox2^CreERT2^ Foxi3cKO mutant incisors typically displayed a completely arrested small bud towards E14.5. However, we also observed that a milder phenotype emerged in some samples: bigger bud or smaller cap. (B) We scored the three different mutant phenotypes and compared WT incisor bud length against each score. Fold change quantifications show a continuous increase in mutant incisor bud length as the phenotype resembled the WT phenotype the most (N=9+3+3+2). (C) Number of samples per score and per litter.

**Figure S2.**
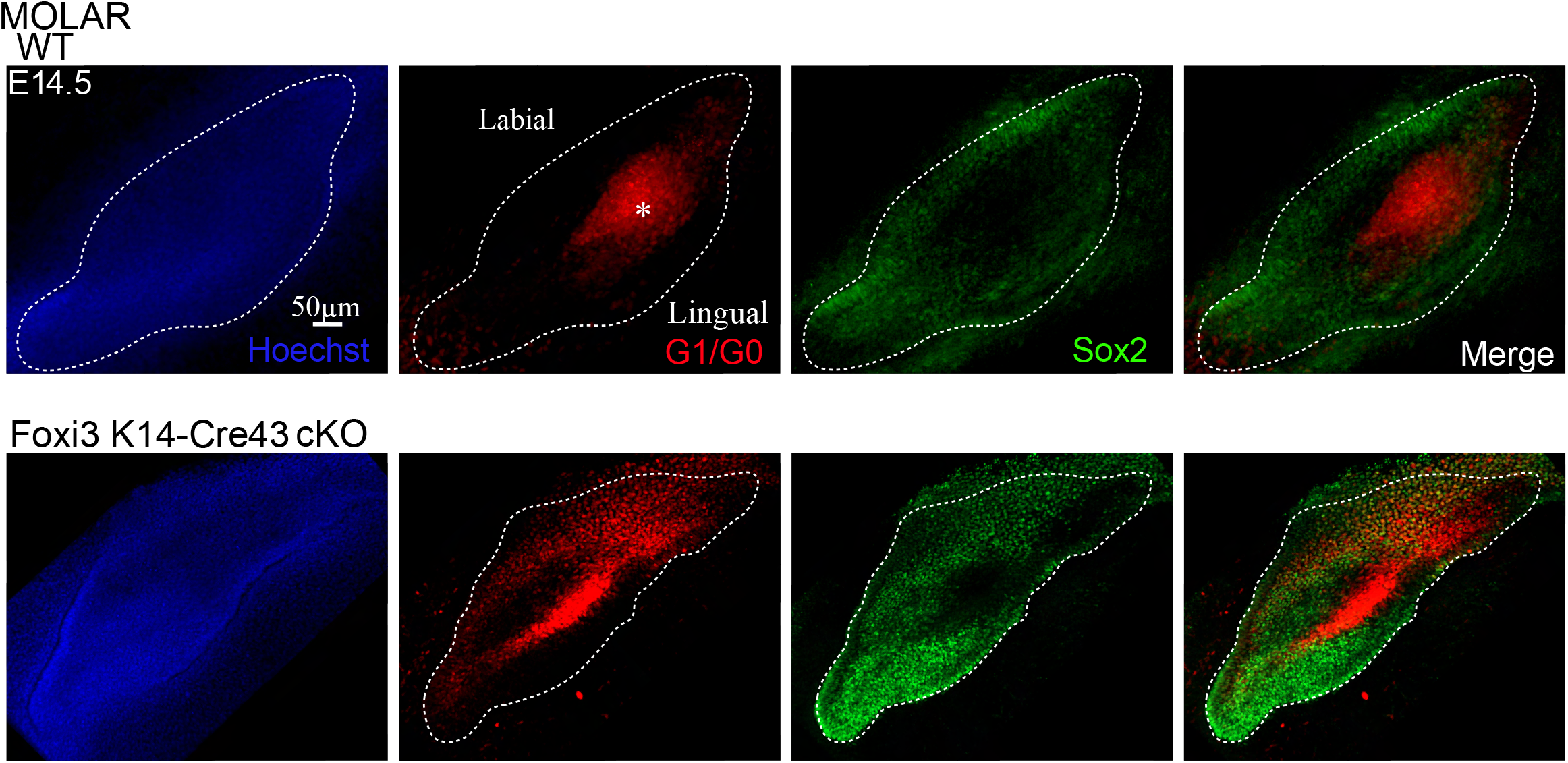
Bud cells abnormally shifted to cell cycle exit do not express Sox2. Sox2 is a marker for pluripotency during development of teeth. Each figure is a volume rendering of the planar xy plane of incisors and molars. Immunofluorescence Sox2 staining in the dental epithelium (green). Fucci G_1_/G_0_ cells (red), and tooth epithelium perimeter (dotted line). Nuclei were stained with Hoechst 33342 (blue). The pEK is marked with an asterisk. In E14.5 WT molars, Sox2 expression is localized surrounding the pEK signaling center region showing an inverse expression pattern. K14^Cre43^ Foxi3cKO mutant molars display the same inverse expression pattern, and a decreased Sox2 expression compared to the WT counterparts due to the increase in G_1_/G_0_ cells distribution.

## Materials and Methods

### Animals

All mouse studies were approved by the National Animal Experiment board. In this study, the following transgenic mouse reporter lines were used: fluorescent cell cycle reporter (Fucci) bitransgenic mice express nuclear red in G_1_/G_0_ phase (Cdt1-mKO) and nuclear green in S/G_2_/M phases (Gem-mAZ) (Sakaue-Sawano et al., 2008); TCF/ Lef:H2B-GFP mice express nuclear GFP as a marker of canonical Wnt/β-catenin signaling activity, which contains several copies of TCF/Lef1 DNA-binding sites driving expression of the H2B-EGFP fusion protein (013752; Jackson Laboratories); and [mK17 5’]-GFP mice which express GFP under the Keratin 17 promoter and visualize the tooth epithelium (023965, Jackson Laboratories).

The following mutant mouse lines were used: the bud stage Foxi3 conditional knock out mutant model was generated by crossing K14^Cre43^; Foxi3^+/−^ males (Andl et al., 2004; Edlund et al., 2014;) with Foxi3^floxed/floxed^ females (Foxi3^tm1.1Akg/J^; a kind gift from Dr. M. Jussila and Dr I. Thesleff) to obtain K14^Cre43^; Foxi3^−/floxed^ mice (Jussila et al., 2015), for analyses from bud stage onwards (E13). For the early stage Foxi3 conditional knock out mutant model (from E11), tamoxifen inducible Sox2^CreERT2^(a kind gift from Dr. F. Michon); Foxi3^+/−^ (a kind gift from Dr. M. Jussila and Dr I. Thesleff) males were crossed with Foxi3^floxed/floxed^ females to obtain Sox2^CreERT^2; Foxi3^−/floxed^ mice. To detect the level of efficiency of the inducible Sox2^CreERT2^ model, ROSA-tDtomato fluorescent inducible reporter female mice (007914, Jackson Laboratories) crossed with Sox2^CreERT2^ males were used to detect the expression of a red fluorescent cytoplasmic protein in cells that express Cre recombinase. Mutant animals were crossed with fluorescent reporters for analysis of cellular behaviors, Foxi3^+/floxed^ animals were used as controls. Embryos were staged according to limb and tooth morphology; vaginal plug day was counted as embryonic day E0.5.

### Tamoxifen administration

A working solution of Tamoxifen (Sigma-Aldrich, T5648) in corn oil (Sigma-Aldrich, 47112-U) was prepared at 50 mg/ml. Tamoxifen solution was sonicated for intervals of 15 × 15 × 7 minutes, and kept at −20°C. To achieve conditional knock down of Foxi3 in the dental epithelium in a time-specific manner, pregnant females were injected twice by intraperitoneal injections with tamoxifen, at E10.5 and E11.5. Tamoxifen daily dose was 2.5mg/30g weigh per mouse together with half the dosage (1.25mg/30g weigh) of progesterone to reduce abortion rates after tamoxifen.

### Tissue preparation and whole-mount immunofluorescence

Embryonic mandibles were dissected at E11.5-E14.5 and whole-mount explants where fixed from 2 h to overnight in 4% PFA. For whole-mount fluorescent staining, fixed tissues were permeabilized with 0.5% TritonX-100 for 2 h RT and washed with PBS. Unspecific staining was blocked by incubation in 5% normal donkey/goat serum, 0.3% BSA, 0.1% TritonX-100 in PBS 1 h RT. Tissues were incubated overnight in 4°C with the primary antibody rat polyclonal anti-mouse CD326 (EpCam, 1:1000, Pharmingen), rabbit polyclonal cleaved caspase 3 (1:400, Cell Signaling Technologies), goat polyclonal Sox2 (1:100; Santa Cruz Biotechnology, Inc.),, and detected with Alexa Fluor-488, Alexa Fluor-568 or Alexa Fluor-647 conjugated secondary antibodies (1:500, BD and Invitrogen) and nuclei were stained with Hoechst 33342. Tissues were mounted with Vectashield (Vector Laboratories).

### Fluorescence confocal microscopy

Fixed samples were imaged with Zeiss LSM700 microscope and HC PL APO 10×/0.45 (air) and LD LCI PL APO 25×/0.8 Imm Corr (water, glycerol, oil) objectives. Images were acquired as z stacks at 1-3 μm intervals. All results represent at least three independent experiments, unless specified otherwise.

### Quantitative and statistical analyses of experimental data

Analyses of images and quantitative measurements were performed with Imaris 9.0.1 (Bitplane). Images were processed for presentation with Photoshop CC and Illustrator CC software (Adobe Systems). Statistical analysis and further graphing were done with Prism 6 (GraphPad Software). All measurements were done in three dimensions from whole-mount volume renderings of confocal optical Z-stacks. For quantifying cell density and proliferation, individual cell borders were visualized and traced in 3D with fluorescent reporters in whole-mount tissues; measurements were done as described previously (Mogollón et al., 2021; Trela et al., 2021). Differences between groups of: bud dimensions, cell number, cell density, and caspase-3 presence were assessed with the Mann-Whitney U test. For analysis of TCF/Lef:H2B-GFP signal intensities, the cut off value for high and low expressing cells was adjusted according to overall signal intensity in each sample.

### RNA extraction and qPCR

Tooth embryonic tissue was homogenized for qRT-PCR into TRI Reagent (Invitrogen) and Precellys 24-homogenizer (Bertin Instruments). RNA extraction was done by guanidium thiocyanate-phenol-chloroform method and then purified by RNeasy Plus micro kit (Qiagen GmbH) according to manufacturer’s instructions. Reverse transcription was done using QuantiTect Reverse Transcription Kit (Qiagen). qRT-PCR was performed using AmpliTaq Gold® DNA Polymerase (Thermo Fisher) with Lightcycler 480 (Roche Applied Science). Data were analyzed using Lightcycler 480 analysis software and normalized against GAPDH. Foxi3 gene expression in K14^Cre43^; Foxi43; Foxi3^−/floxed^ (n=3) and Sox2^CreERT2^; Foxi3^−/floxed^ (n=3) mutant teeth was quantified using ΔΔCT method and compared to control (namely: WT) tissue of corresponding litters (n=6). Results are shown as mean±s.d. Statistical analysis was performed using SPSS (IMB). Statistical significance of the difference between mutant and control samples was tested with Mann-Whitney *U*-test. *P*-value of 0.05 was chosen as the significance threshold. Gene expression levels were normalized using GAPDH. Primers are listed in the following table:

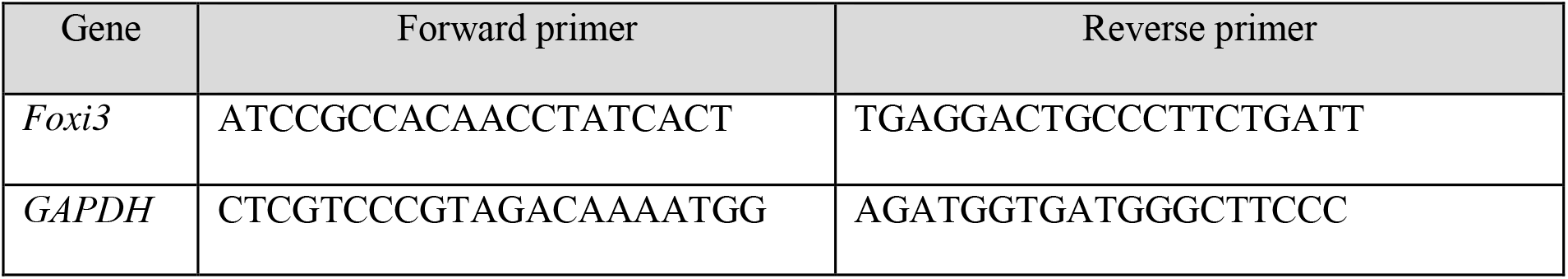

## References

1. Ahtiainen, L., Lefebvre, S., Lindfors, P. H., Renvoisé, E., Shirokova, V., Vartiainen, M. K., Thesleff, I. & Mikkola, M. L. 2014. Directional cell migration, but not proliferation, drives hair placode morphogenesis. Developmental cell, 28, 588–602.

2. Ahtiainen, L., Uski, I., Thesleff, I. & Mikkola, M. L. 2016. Early epithelial signaling center governs tooth budding morphogenesis. Journal of Cell Biology, 214, 753–767.

3. Andl, T., Ahn, K., Kairo, A., Chu, E. Y., Wine-Lee, L., Reddy, S. T., Croft, N. J., Cebra-Thomas, J. A., Metzger, D. & Chambon, P. 2004. Epithelial Bmpr1a regulates differentiation and proliferation in postnatal hair follicles and is essential for tooth development.

4. Benayoun, B. A., Caburet, S. & Veitia, R. A. 2011. Forkhead transcription factors: key players in health and disease. Trends in Genetics, 27, 224–232.

5. Biggs, L. C., Mäkelä, O. J., Myllymäki, S.-M., Roy, R. D., Närhi, K., Pispa, J., Mustonen, T. & Mikkola, M. L. 2018. Hair follicle dermal condensation forms via Fgf20 primed cell cycle exit, cell motility, and aggregation. Elife, 7, e36468.

6. Cobourne, M., Hardcastle, Z. & Sharpe, P. 2001. Sonic hedgehog regulates epithelial proliferation and cell survival in the developing tooth germ. Journal of dental research, 80, 1974–1979.

7. Dassule, H. R., Lewis, P., Bei, M., Maas, R. & Mcmahon, A. P. 2000. Sonic hedgehog regulates growth and morphogenesis of the tooth. Development, 127, 4775–4785.

8. Drögemüller, C., Karlsson, E. K., Hytönen, M. K., Perloski, M., Dolf, G., Sainio, K., Lohi, H., Lindblad-Toh, K. & Leeb, T. 2008. A mutation in hairless dogs implicates FOXI3 in ectodermal development. Science, 321, 1462–1462.

9. Edlund, R. K., Ohyama, T., Kantarci, H., Riley, B. B. & Groves, A. K. 2014. Foxi transcription factors promote pharyngeal arch development by regulating formation of FGF signaling centers. Developmental biology, 390, 1–13.

10. Evans, A. R., Wilson, G. P., Fortelius, M. & Jernvall, J. 2007. High-level similarity of dentitions in carnivorans and rodents. Nature, 445, 78–81.

11. Hallikas, O., Das Roy, R., Christensen, M. M., Renvoise, E., Sulic, A. M. & Jernvall, J. 2021. System-level analyses of keystone genes required for mammalian tooth development. J Exp Zool B Mol Dev Evol, 336, 7–17.

12. Harjunmaa, E., Kallonen, A., Voutilainen, M., HämäLäinen, K., Mikkola, M. L. & Jernvall, J. 2012. On the difficulty of increasing dental complexity. Nature, 483, 324–327.

13. Huh, S.-H., Jones, J., Warchol, M. E. & Ornitz, D. M. 2012. Differentiation of the lateral compartment of the cochlea requires a temporally restricted FGF20 signal. PLoS biology, 10, e1001231.

14. Iurlaro, M., Ficz, G., Oxley, D., Raiber, E.-A., Bachman, M., Booth, M. J., Andrews, S., Balasubramanian, S. & Reik, W. 2013. A screen for hydroxymethylcytosine and formylcytosine binding proteins suggests functions in transcription and chromatin regulation. Genome biology, 14, 1–11.

15. Järvinen, E., Birchmeier, W., Taketo, M. M., Jernvall, J. & Thesleff, I. 2006. Continuous tooth generation in mouse is induced by activated epithelial Wnt/β-catenin signaling. Proceedings of the National Academy of Sciences, 103, 18627–18632.

16. Jernvall, J. & Thesleff, I. 2000. Reiterative signaling and patterning during mammalian tooth morphogenesis. Mechanisms of development, 92, 19–29.

17. Jernvall, J. & Thesleff, I. 2012. Tooth shape formation and tooth renewal: evolving with the same signals. Development, 139, 3487–3497.

18. Jussila, M., Aalto, A. J., Sanz Navarro, M., Shirokova, V., Balic, A., Kallonen, A., Ohyama, T., Groves, A. K., Mikkola, M. L. & Thesleff, I. 2015. Suppression of epithelial differentiation by Foxi3 is essential for molar crown patterning. Development, 142, 3954–3963.

19. Juuri, E., Jussila, M., Seidel, K., Holmes, S., Wu, P., Richman, J., Heikinheimo, K., Chuong, C.-M., Arnold, K. & Hochedlinger, K. 2013. Sox2 marks epithelial competence to generate teeth in mammals and reptiles. Development, 140, 1424–1432.

20. Khatri, S. B., Edlund, R. K. & Groves, A. K. 2014. Foxi3 is necessary for the induction of the chick otic placode in response to FGF signaling. Developmental biology, 391, 158–169.

21. Kupczik, K., Cagan, A., Brauer, S. & Fischer, M. S. 2017. The dental phenotype of hairless dogs with FOXI3 haploinsufficiency. Sci Rep, 7, 5459.

22. Kuony, A. & Michon, F. 2017. Epithelial markers aSMA, Krt14, and Krt19 unveil elements of murine lacrimal gland morphogenesis and maturation. Frontiers in physiology, 8, 739.

23. Lam, E. W.-F., Brosens, J. J., Gomes, A. R. & Koo, C.-Y. 2013. Forkhead box proteins: tuning forks for transcriptional harmony. Nature Reviews Cancer, 13, 482–495.

24. Li, J., Chatzeli, L., Panousopoulou, E., Tucker, A. S. & Green, J. B. 2016. Epithelial stratification and placode invagination are separable functions in early morphogenesis of the molar tooth. Development, 143, 670–681.

25. Lumsden, A. 1988. Spatial organization of the epithelium and the role of neural crest cells in the initiation of the mammalian tooth germ.

26. Matalova, E., Sharpe, P. T., Lakhani, S. A., Roth, K. A., Flavell, R. A., Setkova, J., Misek, I. & Tucker, A. S. 2003. Molar tooth development in caspase-3 deficient mice. International Journal of Developmental Biology, 50, 491–497.

27. Matalova, E., Tucker, A. & Sharpe, P. 2004. Death in the life of a tooth. Journal of dental research, 83, 11–16.

28. Mcgowan, K. M. & Coulombe, P. A. 1998. Onset of keratin 17 expression coincides with the definition of major epithelial lineages during skin development. The Journal of cell biology, 143, 469–486.

29. Mina, M. & Kollar, E. 1987. The induction of odontogenesis in non-dental mesenchyme combined with early murine mandibular arch epithelium. Archives of oral biology, 32, 123–127.

30. Mogollón, I. & Ahtiainen, L. 2020. Live tissue imaging sheds light on cell level events during ectodermal organ development. Frontiers in Physiology, 11, 818.

31. Mogollón, I., Moustakas-Verho, J. E., Niittykoski, M. & Ahtiainen, L. 2021. The initiation knot is a signaling center required for molar tooth development. Development, 148, dev194597.

32. Munne, P. M., Tummers, M., Jarvinen, E., Thesleff, I. & Jernvall, J. 2009. Tinkering with the inductive mesenchyme: Sostdc1 uncovers the role of dental mesenchyme in limiting tooth induction.

33. Mustonen, T., Ilmonen, M., Pummila, M., Kangas, A. T., Laurikkala, J., Jaatinen, R., Pispa, J., Gaide, O., Schneider, P. & Thesleff, I. 2004. Ectodysplasin A1 promotes placodal cell fate during early morphogenesis of ectodermal appendages.

34. Nonomura, K., Yamaguchi, Y., Hamachi, M., Koike, M., Uchiyama, Y., Nakazato, K., Mochizuki, A., Sakaue-Sawano, A., Miyawaki, A. & Yoshida, H. 2013. Local apoptosis modulates early mammalian brain development through the elimination of morphogen-producing cells. Developmental cell, 27, 621–634.

35. Pispa, J., Mustonen, T., Mikkola, M. L., Kangas, A. T., Koppinen, P., Lukinmaa, P. L., Jernvall, J. & Thesleff, I. 2004. Tooth patterning and enamel formation can be manipulated by misexpression of TNF receptor Edar. Developmental dynamics: an official publication of the American Association of Anatomists, 231, 432–440.

36. Pispa, J. & Thesleff, I. 2003. Mechanisms of ectodermal organogenesis. Developmental biology, 262, 195–205.

37. Sakaue-Sawano, A., Kurokawa, H., Morimura, T., Hanyu, A., Hama, H., Osawa, H., Kashiwagi, S., Fukami, K., Miyata, T. & Miyoshi, H. 2008. Visualizing spatiotemporal dynamics of multicellular cell-cycle progression. Cell, 132, 487–498.

38. Sanz-Navarro, M., Seidel, K., Sun, Z., Bertonnier-Brouty, L., Amendt, B. A., Klein, O. D. & Michon, F. 2018. Plasticity within the niche ensures the maintenance of a Sox2+ stem cell population in the mouse incisor. Development, 145, dev155929.

39. Sharir, A. & Klein, O. D. 2016. Watching a deep dive: live imaging provides lessons about tooth invagination. Journal of Cell Biology, 214, 645–647.

40. Shirokova, V., Biggs, L. C., Jussila, M., Ohyama, T., Groves, A. K. & Mikkola, M. L. 2016. Foxi3 deficiency compromises hair follicle stem cell specification and activation. Stem Cells, 34, 1896–1908.

41. Shirokova, V., Jussila, M., Hytönen, M. K., Perälä, N., Drögemüller, C., Leeb, T., Lohi, H., Sainio, K., Thesleff, I. & Mikkola, M. L. 2013. Expression of Foxi3 is regulated by ectodysplasin in skin appendage placodes. Developmental dynamics, 242, 593–603.

42. Trela, E., Lan, Q., Myllymäki, S.-M., Villeneuve, C., Lindström, R., Kumar, V., Wickström, S. A. & Mikkola, M. L. 2021. Cell influx and contractile actomyosin force drive mammary bud growth and invagination. Journal of Cell Biology, 220, e202008062.

43. Tummers, M. & Thesleff, I. 2009. The Importance of Signal Pathway Modulation in all Aspects of Tooth Development. Journal of Experimental Zoology Part B-Molecular and Developmental Evolution, 312b, 309–319.

44. Vaahtokari, A., Aberg, T. & Thesleff, I. 1996. Apoptosis in the developing tooth: association with an embryonic signaling center and suppression by EGF and FGF-4. Development, 122, 121–129.

45. Van Genderen, C., Okamura, R. M., Farinas, I., Quo, R.-G., Parslow, T. G., Bruhn, L. & Grosschedl, R. 1994. Development of several organs that require inductive epithelial-mesenchymal interactions is impaired in LEF-1-deficient mice. Genes & development, 8, 2691–2703.

